# Simple case of prebiotic evolution: vesicle populations can respond to selection for greater turbidity via emergent cooperative dynamics

**DOI:** 10.64898/2026.01.09.698687

**Authors:** Tymofii Sokolskyi, David Baum

## Abstract

Adaptive evolution has long been hypothesized to be possible in the absence of genetic molecules, but experimental evidence remains lacking. Fatty acid vesicles represent an intriguing model to study the emergence of prebiotic evolution, since they can spontaneously grow and divide and have been hypothesized to be capable of non-genetic inheritance. In this study we conducted multiple experiments to test whether vesicle populations can respond to artificial selection for greater turbidity and, if so, whether that response can be tied to an inheritance-like mechanism. We prepared 96 independent vesicle populations, incubated them for 24 hours and then selected half the populations to propagate into the next generation. The populations to propagate were picked either randomly, representing drift controls, or were the populations with the greatest turbidity, representing selection. Population propagation involved resuspension, transfer into fresh buffer, feeding with an amphiphile stock, and then incubating for the next 24 hours. In three replicate experiments run for at least 10 generations, we observed consistently greater turbidity in selection lineages compared to drift, as well as a reduction in the heritability (i.e., the correlation between parent and offspring turbidities). We conducted additional experiments to evaluate whether this response to selection is caused by a simple carryover effect or reflects cooperative dynamics, where vesicles from the parental generation affect newly formed vesicles in the offspring generation. The response to selection is much lower if we omitted the resuspension step and/or if we did not feed transfers with amphiphiles but instead mixed them we pre-formed vesicles. Combined with imaging and other analyses of the resuspension and feeding process, these results suggest that cooperative vesicle dynamics occur, where a small number of intact vesicles from a parental generation alters the dynamics of new vesicle formation following food addition. Overall, this study represents the first experimental finding of a response to artificial selection in prebiotic chemistry.

## Introduction

If one defines life as a self-sustaining chemical system capable of Darwinian evolution^1^, explaining the origins of life requires that we find chemical systems that are capable of adaptive evolution yet simple enough to emerge spontaneously under prebiotic conditions^2^. One class of models argues that the first evolvers were self-organized amphiphile assemblies (micelles, droplets, or vesicles) that could grow and divide and manifest compositional rather than genetic information encoding^3,4,5,6,7^. Amphiphiles such as long-chain such fatty acid are present on meteorites^8^ and comets^9^, which likely delivered small amounts of amphiphiles to prebiotic oceans^8^, and would have supplemented the spontaneous formation of similar molecules on Earth via hydrothermal processes^10,11^. Thus, it is possible that amphiphile assemblies formed in some geological settings, such as in tectonic fault zones^12^. Since such fatty acid vesicles can grow and divide^13,14^, they would be plausible candidates for the first adaptive evolvers provided they could generate heritable variation on which selection could act.

Spontaneously formed prebiotic vesicles could potentially manifest heritability via two distinct mechanism. The first is through compositional inheritance based on the relative abundances of different vesicle-associated and/or encapsulated molecules, as suggested by the Graded Autocatalysis Replication Domain (GARD) model. The GARD model suggests that amphiphile assemblies could replicate and pass compositional information to the following generations due to mutual catalysis, where assemblies are composed of amphiphile species that promote one another’s assembly^3,4,5,6,7^. The second mechanism of inheritance is through a physical phenomenon called the matrix effect, where a set of pre-existing vesicles provided with an excess of unincorporated food cause the formation of new vesicles that are similar in size to the pre-existing vesicles^15^. This phenomenon has been observed with varying food to template ratios and with mixtures of more than one amphiphile species^16^. A proposed mechanism for the matrix effect is that the new amphiphiles insert rapidly into the outer membrane leaflet of a template vesicle, which causes curvature and eventually formation of a second vesicle of similar size to the template^17^. The two mechanisms may not be completely independent because physical features of vesicles, such as size, shape or lamellarity, can be sensitive to amphiphile composition^18,19,20,21^.

Whether inheritance is based on composition or the matrix effect, we might expect that populations of vesicles subjected to selection for an emergent trait might change over time in the selected direction. The only previous experimental study in this area found weak and inconsistent evidence for heritability of size and Nile Red fluorescence in serially-transferred vesicles composed of decanoic acid and decylamine^22^.

Here, we tested whether populations of vesicles can respond to selection for an emergent trait, turbidity, as measured by light scattering at 400 nm. Turbidity is a multifactorial trait that is sensitive to several vesicle characteristics (radius, shape, lamellarity, and lumen contents)^23^. It is also quick and easy to measure in a plate reader, allowing for large sample sizes. We found a significantly higher turbidity in selected versus unselected (drift) vesicles populations. This response to selection was associated with a loss of heritability, which explains why the divergence between selected and unselected lines only changed over the first few generations. Follow-on experiments suggest that this response to selection cannot entirely be explained by carrying forward bulk composition. The matrix effect, or something like it, provides the most plausible mechanism for the inheritance of vesicle characteristics from one generation to the next. Further work is needed, however, to elucidate the mechanism and see if a more consistent and open-ended response to selection would be seen if a greater diversity of amphiphiles were present.

## Results and Discussion

*Vesicles respond to selection for greater turbidity, but only when resuspending and feeding with amphiphile stock*. To test whether vesicle populations can respond to selection for greater turbidity we compared changes in turbidity between selection and drift lines. Each experiment was conducted in two 96-well plates (Dot Scientific, Burton, MI, USA), one composed of selected lineages and the other of unselected (drift) lineages (**Fig. 1**). After a 24-hour incubation each well’s turbidity, absorbance at 400 nm, was measured by a BiotekSynergy HTX spectrophotometer (Agilent Technologies, Santa Clara, CA, USA). In the selected lineages the 48 wells with highest turbidity were each used to seed two wells in the next generation, while the 48 lowest scoring wells left no descendants. In the drift controls, 48 randomly selected wells replicated and the other 48 did not. For both drift and selection plates, “offspring” wells in each following generations were randomized to control for position effects. In most experiments, this procedure was repeated for 10 generations.

**Fig. 1.**
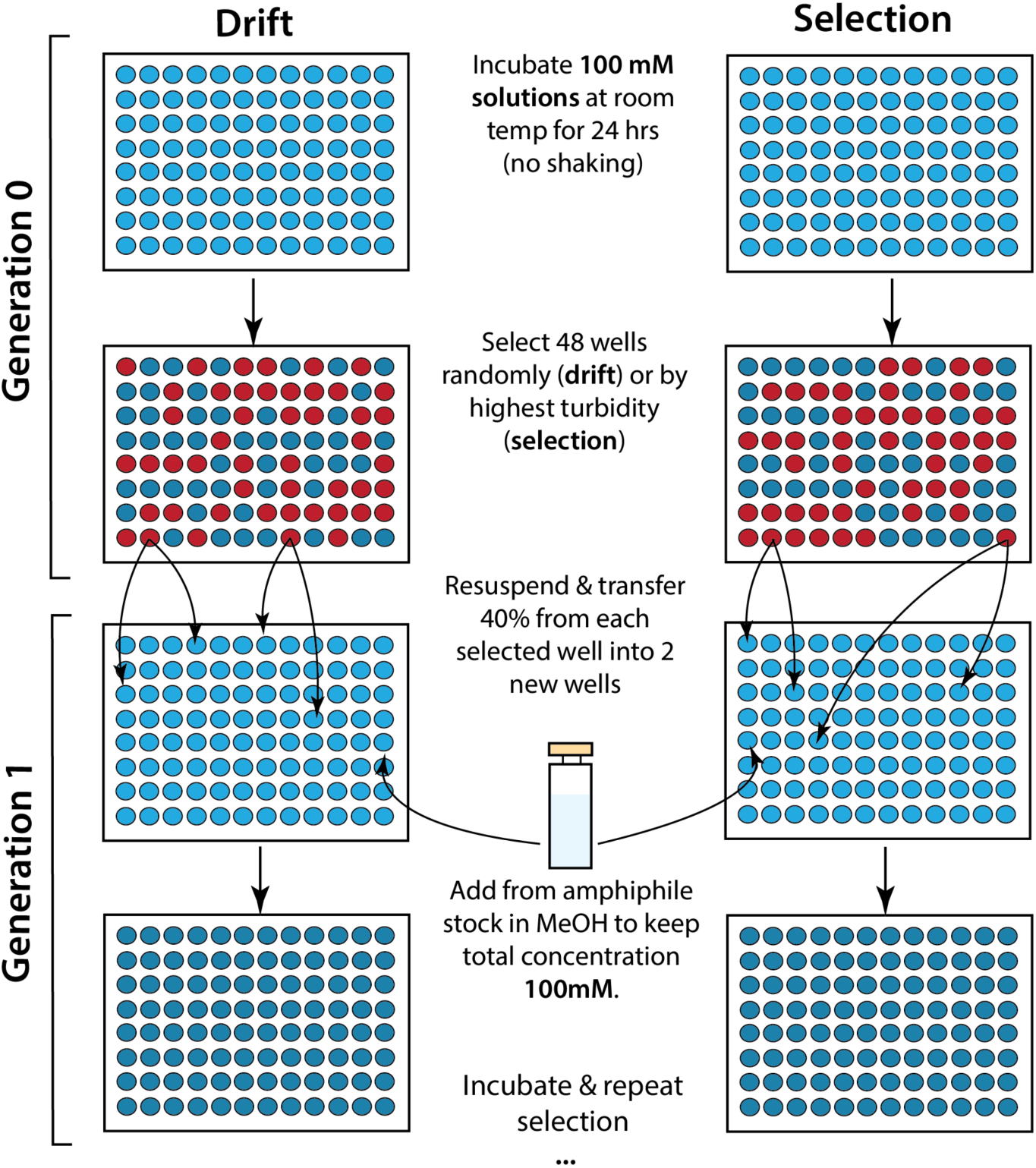
Scheme of selection and drift in the feeding-after-resuspension (FR) protocol.

In the initial experiment the contents of each well were resuspended by repeated (6) rounds of pipetting prior to transferring 40% of the contents to daughter wells containing an appropriate volume of buffer. These offspring wells were then fed with amphiphile monomers, dissolved in methanol, to bring the well’s total amphiphile concentration back to its original concentration of 100 mM. Since this transfer protocol involved feeding after resuspension, we denote it ‘FR.’

Three replicate FR experiments (one run for 20 instead of 10 generations) were conducted, all of which detected a significant difference in turbidity between selected and drift lineages in most generations (**Fig. 2, A; Fig. S1, A-C**). For example, in the 20-generation experiment, denoted FR1 (**Fig. S1, A**), we observed an increase in the drift-selection turbidity difference from generations 3 and 10, after which the difference remained generally above that in drift lineages. FR selection lineages showed consistently lower heritability (the slope of the parent on mid-offspring turbidity) than drift (**Fig. 2, E**). These results show that whatever heritability exists at the start of the experiment is diminished in lineages subject to selection, and is not replaced. In the presence of selection, fewer of the original (generation 0) wells survive than under drift indicating that both descendants of a given parent are more likely to survive (i.e., be selected for) in the selected lineage than in the drift lineages (**Fig. S2, A**).

**Fig. 2.**
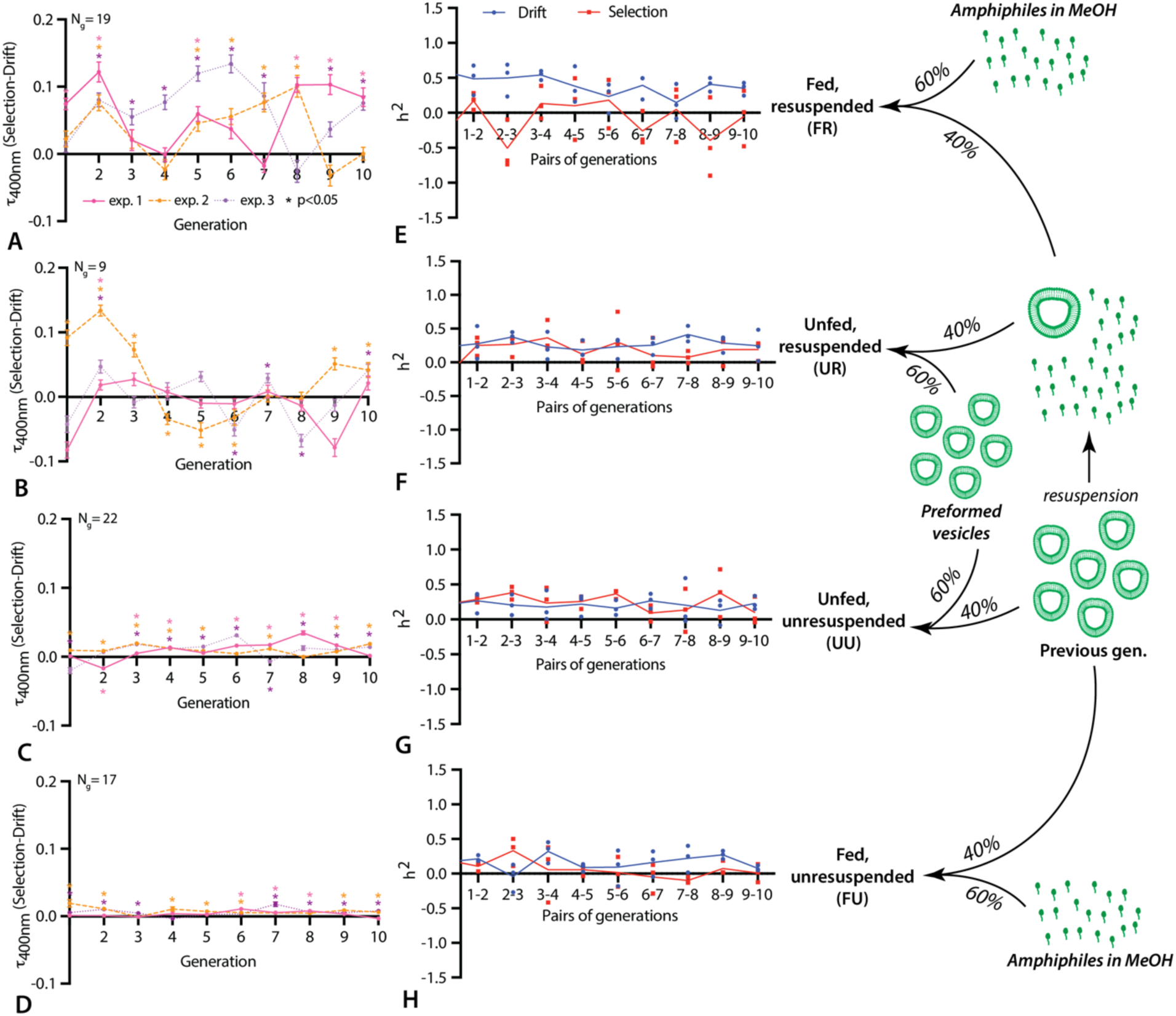
Summary of the results from the selection experiments of four different types. **A, B, C, D** are the selection-drift turbidity difference for the three replicate experiments of the FR, UR, UU and FU protocols, respectively (n=96, error bars are SE). **E, F, G, H** are parent-offspring heritabilities for each pair of generations for each replicate experiment (n=3, lines connect means). N_g_ refers to the total number of generations across the three replicate experiments where selection turbidity is significantly (p<0.05) greater than drift. Asterisks indicate p<0.05 for the selection and drift comparison in **A, B, C, D**.

To see if the response to selection for greater turbidity entails a change in population properties, such as vesicle surface area to volume ratio or the the distribution of different particle types^24,25^, we tracked Nile Red fluorescence (both emission, I_640_, and the degree of blue shift, I_610_/I_660_) in one FR experiment. There was no clear pattern for either metric (**Fig. S3)** and no correlation between turbidity and Nile Red fluorescence for any of the generations (**Fig. S4** and **Table S1**). Although there were statistically significant differences between drift and selection lineages in some generations, the lack of temporal patterns suggests that these reflect day-to-day variation in the experiment or, perhaps, a transfer dependency of Nile Red emission, similar to that observed previously^22^.

### Removing resuspension or feeding with amphiphile stock reduces the response to selection

To see if the formation of new vesicles from the amphiphile stock at the start of each generation was needed for a response to selection we repeated the procedure in triplicate, while replenishing amphiphiles in the form of preformed vesicles. Vesicle solutions were prepared a day before each generation by mixing the methanol amphiphile stock with buffer and incubating for 24 hours in the recipient well. This unfed, resuspended (UR) protocol did not result in any consistent significant turbidity differences between selection and drift lineages (**Fig. 2, B; Fig. S1, D-F**) or in heritability (**Fig. 2, F**). Methanol, the solvent used to introduce amphiphiles cannot explain differences among experimental setups since both stock-fed and vesicle-fed setups contained the same methanol concentration. This suggests that the response to selection in FR transfers depends on new vesicle formation (or growth and division) after the parental vesicles are feed with dissolved amphiphiles.

To evaluate whether resuspension prior to transfer is necessary for the response to selection seen in FR experiments, we conducted a triplicate set of amphiphile-fed experiments that not resuspended before transfer (FU). Here, solutions were removed from parental wells by careful pipetting with a large-bore pipette. In 17 of the 30 generations, turbidity is significantly greater in selected than drift lines, similar to the 19/30 significant values seen in FR (**Fig. 2, D; Fig. S1, J-L**). However, the magnitude of this difference is much smaller and there is no detectable difference in heritability (**Fig. 1, H**) or the number of surviving lineages (**Fig. S2, D**). One possible explanation for the observed significant, but minimal, response to selection is simple carryover, where offspring wells coming from more turbid parental wells are more turbid simply because 40% of their volume came from the parental well. Simple carryover is likely to be more marked in FU that FR because, without resuspension, we would expect a larger fraction of the previous generation’s vesicles to remain intact and contribute to offspring turbidity.

Finally, we conducted a set of vesicle-fed experiments similar to UR, but without resuspension. In this UU setup, where intact vesicles from the parental generation are mixed with pre-formed vesicles, we observed a consistently greater turbidity in selection lineages (**Fig. 2, C; Fig. S1, G-I**). However, as with FU, the magnitude of the turbidity difference was small and heritability did not differ between selection and drift treatments (**Fig. 2, G**), suggesting that the result may reflect carryover only. To evaluate whether vesicle-vesicle interactions might be contributing to the (modest) response to selection in UU experiments, we conducted a fourth experiment for five generations with each transfer done immediately after mixing, without any incubation. The lack of an incubation period in this experiment would presumably prevent interactions between the parental and newly introduced vesicles. This rapid UU procedure yielded a similar response to selection as the 24-hour per generation UU procedure (**Fig. S5**) consistent with the hypothesis that selection-drift differences seen in the UU protocol are due to simple carryover.

Overall, we observed that protocols without resuspension, where intact vesicles are transferred between generation, have a response to selection that is likely only due to simple carryover. The UR protocol showed no response to selection, whereas the FR protocol had the clearest response to turbidity selection and was also the only protocol showing a markedly lower heritability in selected lineages (**Fig. 3, A**). FR is also unique in showing a clear inverse relationship between heritability and mean turbidity (**Fig. 3, B**). These findings suggest the possibility that while the carryover effect is present in all protocols that lacked resuspension, the response to selection seen in the FR protocol reflects an inheritance-like process where the properties of post-resuspension vesicles are somehow transmitted to newly formed vesicles produced from amphiphiles following transfers.

**Fig. 3.**
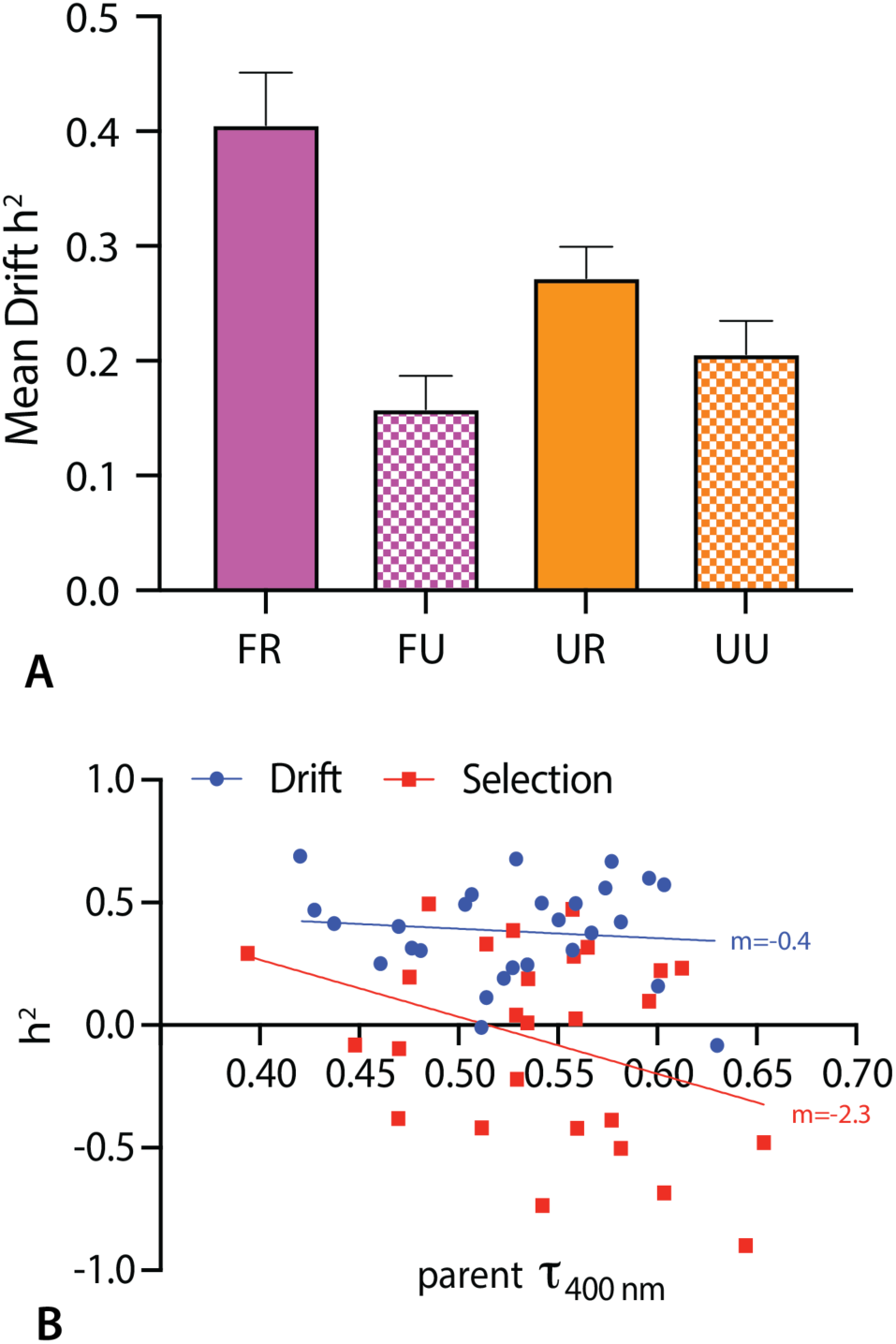
Patterns of heritability changes in different experiments. **A** – mean heritability in drift lineages across all generations of the four experiment types; **B** – linear regression between mean parent generation turbidity and heritability of the respective parent-offspring generation pair across three replicate FR experiments. For **A**, n=30 and error bars are SE, for **B** slope values are indicated on the graph.

### Resuspension induces vesicle reorganization

To elucidate the mechanism underlying the response to selection seen in FR experiments, we used a variety of methods to investigate the transfer and feeding process. Fluorescent microscopy revealed that resuspension greatly reduces the number of large vesicles (**Fig. 4, A-B**), as confirmed by dynamic light-scattering which showed a significant reduction in average vesicle size (**Fig. 4, C**). These changes in the assembly size distribution likely explain the immediately increased turbidity that occurs following resuspension (**Fig. 4, D**).

**Fig. 4.**
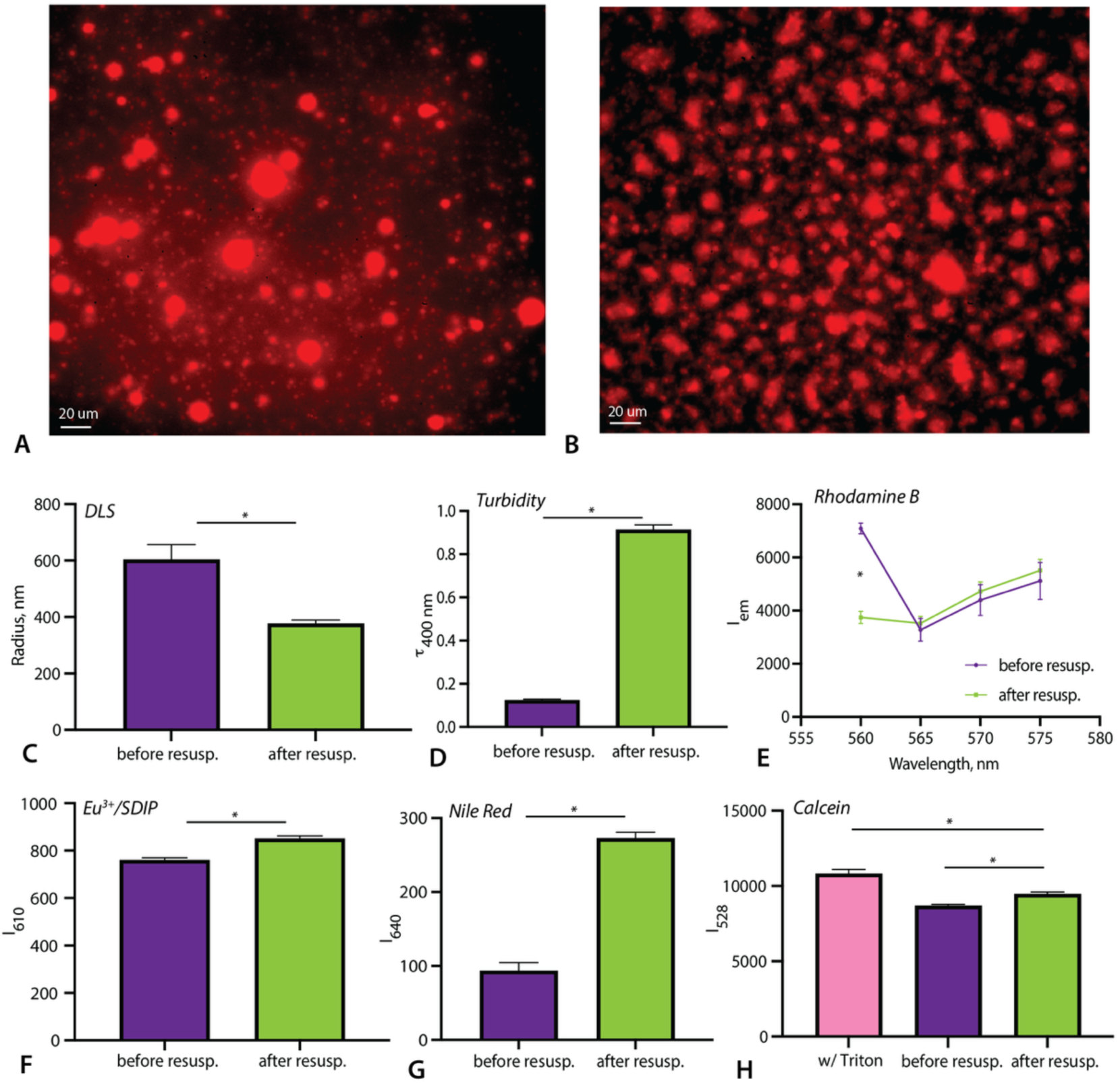
Changes occurring during vesicle resuspension. **A** – Microscopy image of rhodamine-stained vesicles after 24 hours of incubation (before resuspension). **B** – Microscopy image of rhodamine-stained vesicles after 24 hours of incubation and then resuspension: **C-G** – physical changes in vesicle populations before and after resuspension: average vesicle radius (**C**; n=8); turbidity at 400 nm (**D**, n=96); rhodamine B fluorescence at 545 nm excitation (**E**, n=24); SDIP/Eu^3+^ fluorescence at 610 nm (**F**, n=3); Nile Red fluorescence at 640 nm (**F**, n=8). **H** – calcein fluorescence at 528 nm before and after resuspension and a positive control containing 1% Triton X-100, n=10. Error bars are SE. Asterisks indicate comparisons that are significant with a Student’s t-test (p<0.05).

To evaluate whether resuspension disrupts vesicles, we measured Rhodamine B fluorescence which is known to diminish when rhodamine escapes into an aqueous environment^26^, corroborated by a test of rhodamine B fluorescence in different solvents without amphiphiles (**Fig. S6**). Rhodamine fluorescence in vesicle solutions was lower after resuspending, which supports the hypothesis that vesicle membranes had been damaged (**Fig. 4, E**). We also used the Eu^3+^/SDIP chelating assay, developed by Biotium (Fremont, CA, USA), with SDIP added in the input population and europium provided in the dilution buffer added after transfer, which revealed an increase in fluorescence following resuspension (**Fig. 4, F**). Other evidence of vesicle disruption during resuspension comes from higher Nile Red emission at 640 nm, which suggests that resuspending causes an increase in vesicle area to volume ratio (**Fig. 4, G**).

We also tracked vesicle size over the course of incubations in a batch of generation 0 and 1 using DLS. Over the course of a 24 incubation (without any prior generations), vesicle radius increases slightly, but not significantly (p=0.13; **Fig S9, A**), during a 24 hour incubation. Over the generation 1 FR transfer and incubation, size increases more considerably up to 900 nm (**Fig. S9, B**). Note, however, these experiments were done in 1 mL volume in cuvettes, so these results may not be comparable to the FR experiments in 200 µL 96-well plates.

We observed that calcein fluorescence in prepared vesicle solutions declined immediately after resuspension and continued to decline over the next 3 hrs (**Fig. 4, H; Fig. S7-8**). This likely indicates that leakage and/or breaking of the vesicles is induced by resuspension. It should be noted, however, that removing unencapsulated calcein in this analysis is impossible because size-exclusion chromatography affects our vesicles and increases turbidity in a similar manner to resuspension. However, since presence of the vesicle-disrupting detergent Triton X-100 causes an increase in calcein fluorescence (**Fig. 4, H**), we can conclude that there is still a fraction of intact vesicles left after resuspension. A similar pattern to these generation 0 results is seen in the timeseries of generation 1, following a FR transfer, although the equilibrium state is achieved in a slightly longer time (**Fig. S8**).

### A response to selection depends on a period of interaction between parental material and dissolved amphiphiles

Except for the UU protocol, turbidity was dynamic over the course of the first incubation, showing a transient increase and then a steady decline (**Fig. S10**). The FR and FU protocols showed a marked peak of turbidity, both within the first hour, absent when the amphiphile stock solution was not added (**Fig. S11**). In FR protocol, which combines resuspension and methanol stock feeding, there was a second increase in turbidity between 2 and 6 hours that is absent from the other protocols. This led us to suspect that there may be a cooperative process resulting from vesicle formation or interaction with the food in that time frame that could explain the stronger response to selection seen in FRs.

To test this hypothesis, we conducted multiple FR selection experiments with varying generation times (**Fig. 5**). With transfer every 0.5 to 1.5 hrs we observed no drift-selection turbidity difference and overall turbidity decreased after the first 1-3 generations (**Fig. 5, A-B**). This indicates that, at those timepoints, the amphiphile stock does not contribute enough new assemblies to prevent serial dilution of the initial turbidity (the initial generation was preceded by a 24-hour incubation). With 5 hours incubations, there was no turbidity decrease and 3 out 5 generations showed elevated turbidity in selected populations (**Fig. 5, C**), similar to that seen in the 24-hour transfer (**Fig. 5, D**). This confirms that a heritable process contributing to the response to selection begins between 1.5 and 5 hours. This time window corresponds to an almost linear increase in turbidity (**Fig. 5, E**), which is absent in timecourses without amphiphile stock addition (**Fig. S11**). Since turbidity scales with the logarithm of vesicle size, this linear increase is probably indicative of increases in vesicle concentration rather than vesicle size^23^.

**Fig. 5.**
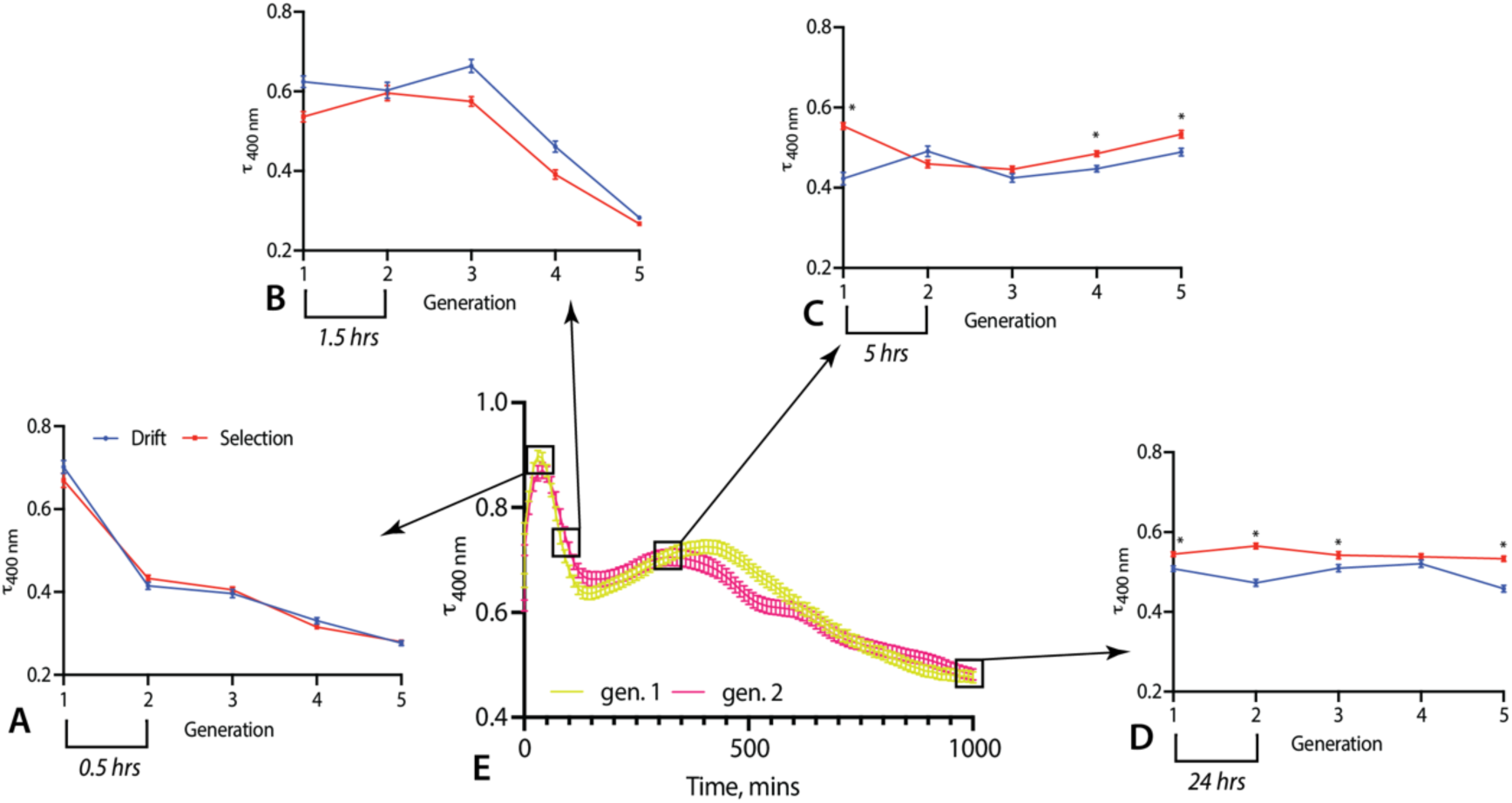
FR selection experiments with varying length of incubations: **A** – 0.5 hrs, **B** – 1.5 hrs, **C** – 5 hrs, **D** – 24 hrs (the latter is derived from the average of the three replicate experiments shown in in Fig. 2**, A**). **E** – time course of turbidity change in the first two generations of an FR transfer, with time-points used for the 5-generation transfer experiments boxed. Asterisks indicate p<0.05 between selection and drift. n=96, error bars are SE.

### Emergent vesicle dynamics supports a non-trivial response to selection but does not enable open-ended evolution

Considering the persistence of some transfer vesicles through resuspension and the high heritability seen at the start of the FR experiments, there are 4 plausible processes that may be occurring following introduction of the amphiphile stock: 1) cooperative vesicle assembly where de novo vesicle formation from food molecules is sensitive to characteristics (such as size) of the preexisting vesicles; 2) independent assembly of food into vesicles followed by spontaneous fusion/fission interactions with the transfer vesicles;; 3) growth of transferred and newly formed vesicles via monomer or micelle incorporation; 4) breakdown of transfer vesicles and assembly of new vesicles from pooled food amphiphiles. Based on the lack of response to selection seen with 1.5-hour incubations but positive response seen with 5-hour incubations, heritability must be based on processes that begin after 1.5 hours of incubation. The linearity of the turbidity increase between 1.5 and 6 hours suggests that de novo vesicle formation is the most likely process, however, this must be verified with additional studies. Prior to this timepoint, the rapid increase and then decrease of turbidity likely corresponds to the formation and dissolution, respectively, of non-vesicle aggregates. This could be an explanation for decrease in turbidity seen with 30-minute incubations (**Fig. 5, A**).

Due to the large volumes required for DLS we were unable to directly track vesicle size over the course of each experiment. However, Nile Red peak emission data (**Fig. S3, A**) suggests that it is unlikely that, after generation 0, vesicle size changes significantly over the course of the FR experiments.

In FR, FU and UU transfer protocols we see higher turbidity in selection lineages compared to drift controls. There was an increase in turbidity in FR experiments, which is similar to responses to artificial selection seen in biology, where heritability is lost following selection^27^. The failure of heritability to decline in FU and UU experiments, suggests that the response to selection in these cases may have a difference mechanism that FR.

In the FU protocol, many generations show increased selection turbidity (17 compared to 19 in FR; **Fig. 2, D**), but the absolute difference between selection and drift heritability and mean drift heritability is the lowest of all experiments (**Fig. 3, A**). This protocol allows a large population of vesicles from the parent to influence the formation of new vesicles forming from dissolved amphiphiles. One might expect, therefore, a stronger response to selection than was seen with the FR protocol. However, an inverse relationship between template vesicle concentration and heritability of vesicle size was observed previously for the matrix effect^17^. This aligns with a proposed mechanism for the effect, which requires overloading of the outer leaflet of template vesicles due when there is a large excess of unincorporated amphiphiles, which is proposed to lead to a doubling in vesicle radius followed by division into two equal vesicle, resulting in the “inheritance” of vesicle size. This could explain the stronger response to selection seen in FR than FU experiments, which is consistent with the matrix effect, or something like it under the FR protocol.

In UR and UU transfers, 60% of the solutions are vesicles prepared independently every generation. Therefore, there is limited potential for a parental sample to influence the next generation in any way other than simple carryover. These experiments thus provide a baseline selection-drift turbidity difference when compared to their respective stock-fed experiments. In UR experiments there is no clear evidence of a response to selection: 9 generations have significantly greater selection turbidity and 4 are significantly smaller (**Fig. 2, B**). The apparent selection/drift difference in experiments UU1-3 (**Fig. 2, C**) is most likely explained by carryover. Although it is conceivable that fusion-fission dynamics occurring when two populations of vesicles are mixed could induce some form of inheritance, this does not seem to be a factor here, since we observe a similar response to selection when the generation time is reduced to 30 min (**Fig. S5**). This would make vesicle-vesicle interactions extremely unlikely, especially since turbidity does not change during incubation in the UU setup (**Fig. S10**). Taken together, these data suggest that observed selection-drift turbidity differences in unresuspended experiments are the result of simple carryover rather than being due to true inheritance of vesicle properties from parental to offspring generations.

Comparing the results from all experiments, we can confidently claim, while the response to selection seen in UU and FU is largely a trivial outcome of vesicle carryover, the response seen in the FR is due to inheritance-like dynamics, likely involving vesicle growth and division and/or new vesicle formation. This heritability is apparently stronger after vesicle populations are resuspended such that a lower fraction of intact vesicles are transferred, which is similar to what was observed for the matrix effect^17^, suggesting that a similar phenomenon occurs in the FR protocol.

## Conclusions

These results are the first definitive evidence that lipid vesicles can respond to selection for their physical properties in a similar manner to extant biology, but without genetic molecules. However, the response to selection seen was immediate and short-lived. There is no evidence, even after 20 generations, that new heritable variation emerges, suggesting a lack of the open-endedness that characterizes biological evolution. Nonetheless, it is striking that the FR experiments demonstrate non-trivial heritability, where information passes from parental to offspring generations by means other than simple carryover.

To better understand the mechanistic basis for the response to selection in FR experiments future work could involve molecular dynamics simulations. Experimentally, it would be desirable to make real-time observations of vesicles that have been resuspended when few with new amphiphiles. Additionally, it would be interesting to target other amphiphile species or utilize different feeding strategies, for example supplying micelles instead of amphiphiles dissolved in methanol, to determine whether non-trivial inheritance is sensitive to these factors.

The mechanism for observed response to selection appears to involve physical interactions among vesicles rather than the compositional inheritance assumed by the GARD model. As our experiments only include 2 amphiphile species (excluding ionic states), the possibilities of compositional inheritance seem limited. However, our experimental framework could be readily deployed in cases where there is a much greater diversity of amphiphiles present to see if a more open-ended response to selection is seen. If so, this result would go a long way to supporting the hypothesis that compositional inheritance could have played an important role in prebiotic evolution prior to the acquisition of nucleic acid polymers.

## Acknowledgements

The authors would like to thank Zoe Todd, Betül Kaçar, John Yin and Annie Bauer for insightful feedback on this study, Anushka Malaiya and Walid Lakdari for assistance with preliminary experiments, and the labs of Jo Handelsman and John Denu for providing access to the plate spectrophotometers. The authors gratefully acknowledge the use of facilities and instrumentation as supported by NSF through the Wisconsin Materials Research Science and Engineering Center (DMR-1720415) and Anna Kiyanova and Mike Efremov from the Soft Material Characterization Lab for technical support. TS was supported by a National Aeronautic and Space Administration Future Investigators in NASA Earth Science and Technology award (80NSSC24K1810). Research funds we provided by the National Science Foundation (grant number DEB 2218817), a Rosemary Grant Advanced Award from the Society for the Study of Evolution, and the University of Wisconsin-Madison Department of Botany.

## Experimental section

### Vesicle preparation

For vesicle stock preparation we used decanoic acid or DA (Pfaltz & Bauer, Waterbury, CT, USA), 1-decanol or DOH (Santa Cruz Biotechnology, Dallas, TX, USA), and methanol (Fischer Scientific, Hampton, NH, USA). For the preparation of the 0.1M pH 7 phosphate buffer (PB) we used monobasic sodium phosphate monohydrate (Millipore Sigma, Burlington, MA, USA) and anhydrous dibasic sodium phosphate (Dot Scientific, Burton, MI, USA). All pipetting of solutions containing vesicles was done with a p1000 pipette since unpublished preliminary experiments showed that smaller pipette tips disrupted vesicle integrity. Since different pipetting styles can affect the turbidity of vesicle populations, all transfers were done by the same person, the lead author.

### Experimental setup

To test whether vesicle populations can respond to selection for greater turbidity we compared changes in turbidity over generations with and without selection. Each experiment was conducted in two 96-well plates (Dot Scientific, Burton, MI, USA), one composed of selected lineages and the other of unselected (drift) lineages. In selected lineages the 48 wells with highest turbidity were allowed to replicate (each was used to seed two wells in the next generation), while the 48 lowest scoring wells each left no descendants. In the drift controls, 48 randomly selected wells replicated.

The total volume in each well was always 200 µL. For all experiments, generation 0 (the generation before the first transfer), was initiated by the addition of 180 µL of PB to each well, followed by 20 µL of a 1M 2:1 DA:DOH stock solution (in methanol). Then, each well was resuspended with a p1000 pipette 5 times and plates were sealed with a plastic seal (Thermo Fischer Scientific, Waltham, MA, USA), taped around the edges to prevent evaporation, and left on a bench at room temperature for 24 hours. In all subsequent generations, 48 of the 96 wells on a plate were transferred into two wells of a new plate. For both drift and selection plates, “receiver” wells were randomized to control for position effects. At the end of each generation, just prior to transfers, turbidity was measured by a Biotek Synergy HTX (Agilent Technologies, Santa Clara, CA, USA) spectrophotometer as absorbance at 400 nm.

The same selection and drift experiments were conducted using four alternative protocols, which we call: fed & resuspended (FR), fed & unresuspended (FU), unfed & resuspended (UR), and unfed & unresuspended (UU). In FR experiments, food, in the form of amphiphile monomers in methanol, was added to vesicles from the previous generation (**Fig. 1**). Each well of a plate from a prior generation was resuspended 5 times with a p1000 pipette and then 80 µL was transferred into 108 µL of PB in a receiver well on a new plate. Then, 12 µL of the 1M 2:1 DA:DOH stock in methanol was added to each well on the new plate. We conducted 3 experiments of this type with an almost identical setup – FR1-3. FR1 lasted for 20 generations, while FR2 and FR3 ended after 10 generations. For FR3 we included 0.2 mM of Nile Red (Neta Scientific, Marlton, NJ, USA) in the amphiphile stock. Three replicate FU experiments were conducted in a similar manner except that the transfer population was not resuspended but was instead transferred with gentle pipetting using a p1000 pipette tip.

For UR transfers, resuspended transfer material from a previous generation was added to wells that contained pre-formed vesicles. A day before each transfer, 108 µL of PB was mixed with 12 µL of the 1M amphiphile stock in each well of the receiver plate, resuspended 5 times. The plate was then sealed and left for 24 hours to form vesicles. Each transferred well from the previous generation was resuspended five times and 80 µL was transferred into the prepared receiver plate. We conducted three experiments of this type, UR1-3, each for 10 generations.

For UU experiments we prepared a vesicle food solution for each well a day before the transfer, as described for UR. However, in UU experiments transfers involved gentle pipetting to keep vesicles as intact as possible. We conducted three replicate experiments of this type, UU1-3, each for 10 generations. UU1 and UU2 were done with the same generation 0 setup as FR1-3 and UR1-3, however, for UU3 we started with a generation 0 that received one 40% transfer and thus had similar turbidity to the values of FR1-3. We also conducted 5 generation transfer experiment, UU4, which omitted the usual 24-hour incubation. Instead, transfers were conducted sequentially using food vesicles prepared the day before.

### Calcein fluorescence

To determine the degree of vesicle leakage during the course of a 24 incubation and during pipetting between generations, we used calcein as a fluorescent probe. A 5 mM stock solution of calcein was prepared by dissolving the powder (Santa Cruz Biotechnology, Dallas, TX, USA) in methanol. This solution was added to PB to a final calcein concentration of 0.125 mM. The resulting solution was dispensed into 20 wells of a black 96-well plate and a 2:1 DA:DOH stock in methanol was added to a final concentration of 100 mM. Emission at 528 nm following excitation at 485 nm was measured every 30 mins using the Biotek Synergy HTX (Agilent Technologies, Santa Clara, CA, USA) spectrophotometer. To prepare the positive control – a solution without vesicles but the same amphiphile concentrations – we added 1% of Triton X-100 (Santa Cruz Biotechnology, Dallas, TX, USA) to the calcein buffer and resuspended the wells after amphiphiles were added. It is known that 1% Triton is sufficient to prevent vesicle formation in these conditions^22^. After 24 hours, 40% of each of sample was transferred into fresh PB (lacking calcein) in a new black plate and the methanol-amphiphile stock solution was added to a final concentration of 60 mM of 2:1 DA:DOH. Fluorescence was also tracked every 30 mins. For each analysis, fluorescence of vesicle samples was divided by the fluorescence of calcein buffer to account for a reduction in fluorescence over the period of incubation and transfer.

### Nile Red

Using a BioTek Synergy HT4 microplate spectrophotometer (Agilent Technologies, Santa Clara, CA, USA) we tracked emission at 610, 640 and 660 nm following excitation at 480 nm every generation. Intensity of fluorescence at 640 nm is proportional to surface area to volume ratio of the particle stained by Nile Red^24^. Meanwhile ratio of intensity at 610 to 660 nm, or the blue shift of the Nile Red emission spectrum, is an indicator of micropolarity of the system^25^.

### Test to assess vesicle disruption during resuspension

Fluorescence-based assays using Rhodamine B and SDIP/EU^3+^ were with a BioTek Synergy HT4 microplate spectrophotometer using 200 μL samples containing 100 mM amphiphiles. Rhodamine B (Neta Scientific, Marlton, NJ, USA) was prepared at 6 μM concentration in a 1M amphiphile methanol stock. Its emission was determined following excitation at 545 nm before and after resuspension. To introduce SDIP into vesicles, 1M amphiphile methanol stock with 20 mM SDIP (Biotium, Fremont, CA, USA) was allowed to incubate for 24 hours in microplates. Then 40% of this vesicle population was transferred into PB containing 0.2 mM EuCl_3_ (Biotium, Fremont, CA, USA). Before and after resuspension of this mixture, emission of the Eu^3+^/SDIP complex was collected at 610 nm following excitation at 280 nm.

### Dynamic Light Scattering

To test quantify vesicle radius we conducted a short transfer experiment in cuvettes. For generation 0, 100 mM DA:DOH vesicles were prepared using 100 µL of the 1M methanol stock and 900 µL of PB in 8 replicate cuvettes. They were measured at 0 and 24 hours and again following transfer of 400 µL, with resuspension using a p1000 pipette, into new cuvettes containing 540 µL of PB. Then 60 µL of the amphiphile stock in methanol was added and the resulting generation 1 samples, which were measured at 0 and 24 hours as well. To obtain Z-average radii, dynamic light scattering (DLS) measurements were done with a Zetasizer Nano (Malvern Pananalytical, Malvern, UK) with 175° backscattering at 25 °C.

