## Supplementary material for "Simple case of prebiotic evolution: vesicle populations can respond to selection for greater turbidity via emergent cooperative dynamics": Fig. S

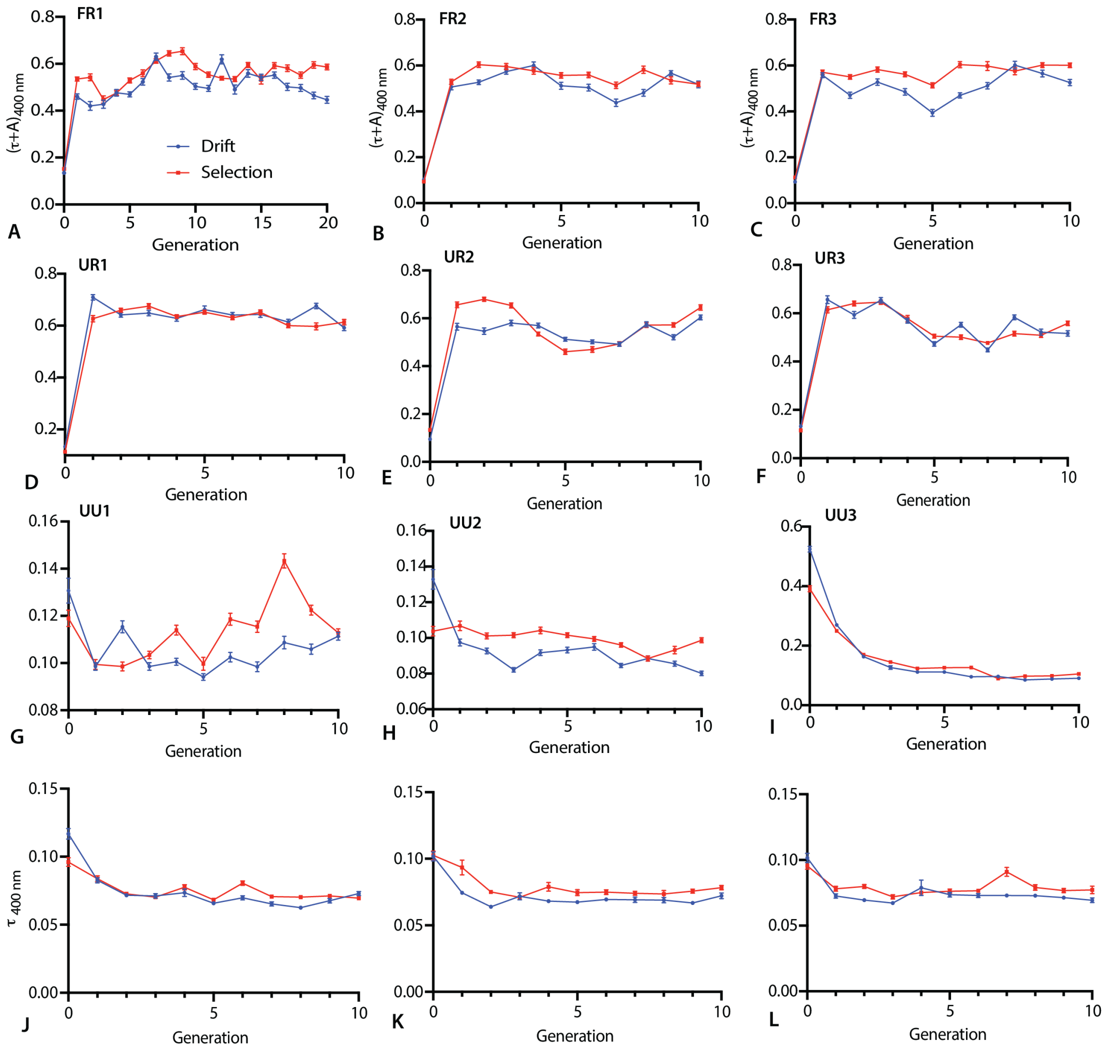


**Fig. S1.** Raw results of the replicate experiments: **A, B, C** – fed, resuspended transfers 1, 2, 3; **D, E, F** – unfed, resuspended transfers 1, 2, 3; **G, H, I** – unfed, unresuspended transfers 1, 2, 3; **J, K, L** – fed, unresuspended transfers 1, 2, 3. n=96, error bars are SE.

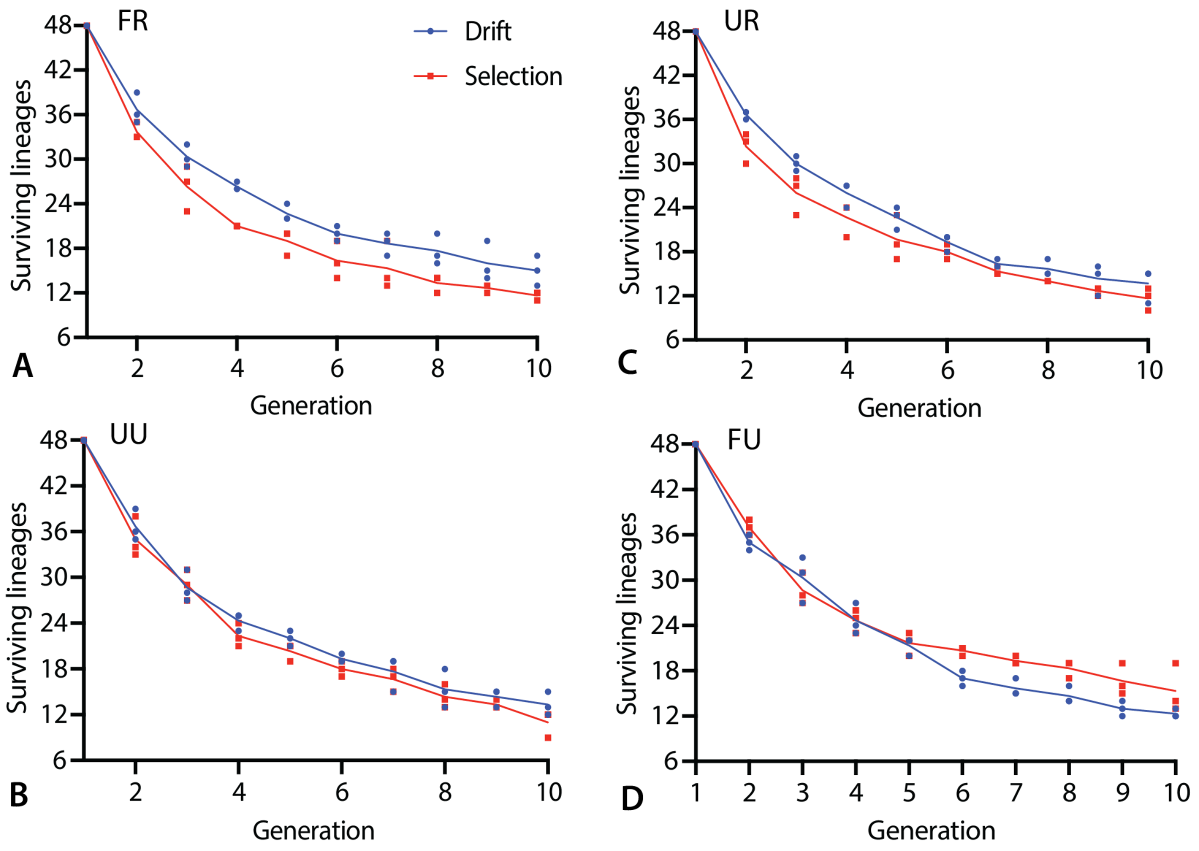


**Fig. S2.** Number of surviving descendants of the original generation 0 lineages for drift and selection plates in each experiment type. **A** – fed, resuspended transfers 1, 2, 3; **B** – unfed, resuspended transfers 1, 2, 3; **C** – unfed, unresuspended transfers 1, 2, 3; **D** – fed, unresuspended transfers 1, 2, 3. n=3, lines connect means.

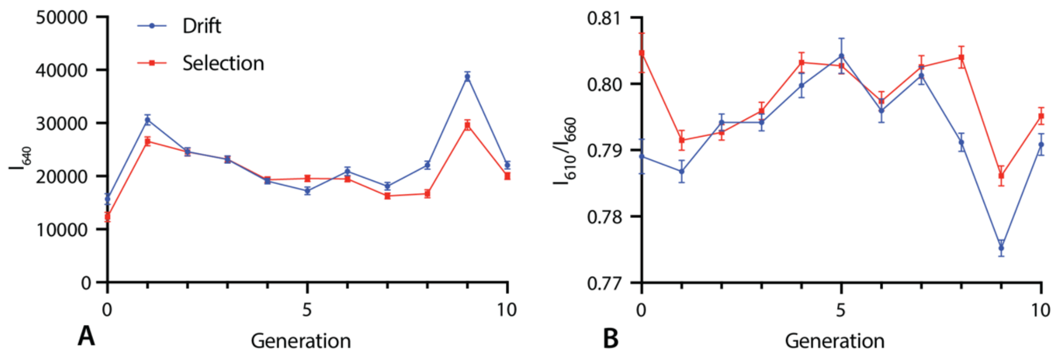


**Fig. S3.** Nile Red fluorescence measurements for FR3: **A** – peak emission at 640 nm; **B** – ratio of emission at 610 nm to 660 nm. n=96, error bars are SE.

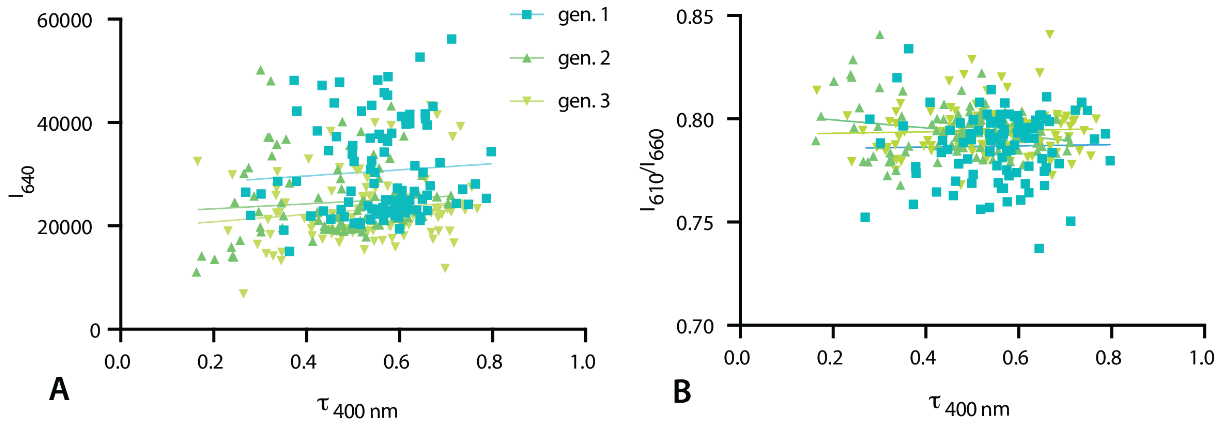


**Fig. S4.** Correlation plots between turbidity and Nile Red emission for first 3 generations of drift lineages for FR3: **A** – peak emission at 640 nm; **B** – ratio of emission at 610 nm to 660 nm. n=96.

| Generation | Drift, I640 | Selection, I640 | Drift, I610/660 | Selection, I610/660 |
|---|---|---|---|---|
| 0 | 0.25610891 | 0.33371921 | -0.0267633 | -0.01639 |
| 1 | 0.09376654 | 0.01366727 | 0.04197177 | 0.03171389 |
| 2 | -0.1114317 | -0.0040383 | -0.0588242 | -0.0523626 |
| 3 | -0.0727198 | -0.0928299 | 0.03872105 | -0.1037585 |
| 4 | 0.02313645 | 0.01001304 | -0.1365998 | -0.0196845 |
| 5 | -0.062271 | -0.073922 | 0.03145417 | -0.1433735 |
| 6 | -0.0187136 | -0.1508559 | 0.03986962 | -0.1040564 |
| 7 | -0.0846738 | -0.1263727 | -0.0942669 | 0.0404701 |
| 8 | -0.025657 | 0.05980209 | -0.1410367 | -0.0585042 |
| 9 | -0.1025235 | -0.0859262 | -0.013623 | 0.23455298 |
| 10 | -0.1714972 | -0.156045 | 0.05493823 | 0.09855872 |

**Table S1.** Pearson correlation coefficients between turbidity and either peak Nile Red emission at 640 nm or the ratio between emission at 610 nm and 660 nm for both selection and drift lineages in FR3.

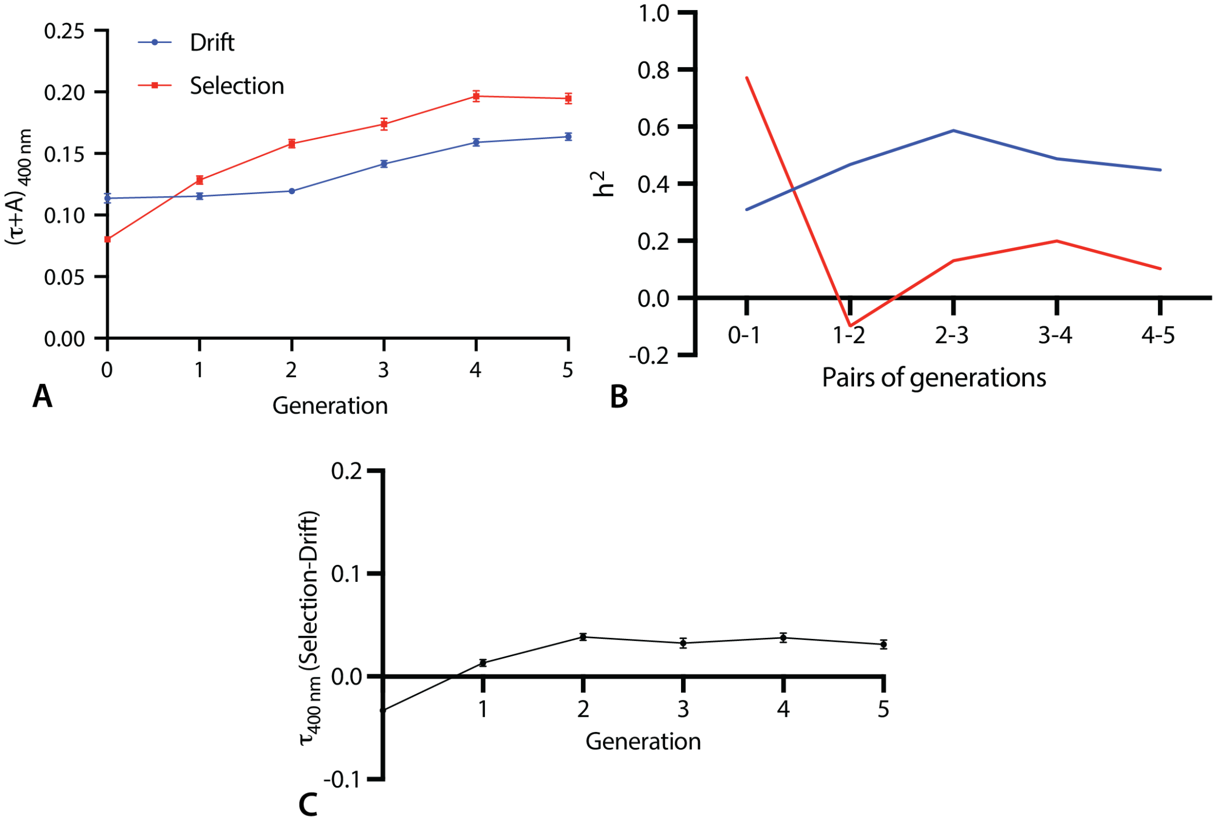


**Fig. S5.** Turbidity changes (A) and heritability (B) in the short-term unfed transfer experiment UU4.

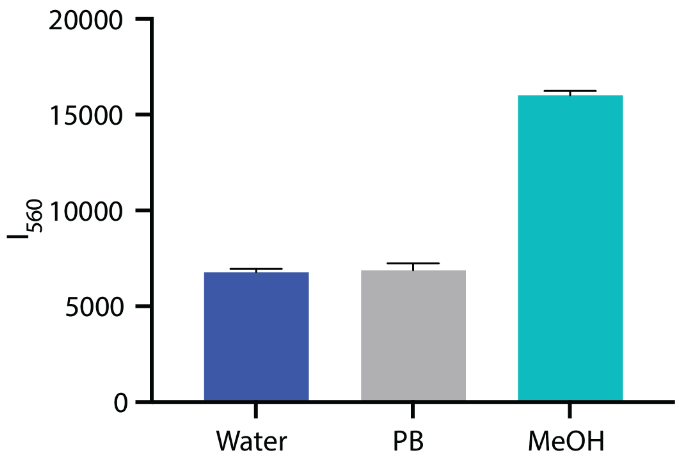


**Fig. S6.** Rhodamine B emission at 560 nm following excitation at 545 nm in different solvents without amphiphiles. n=6, error bars are SE.

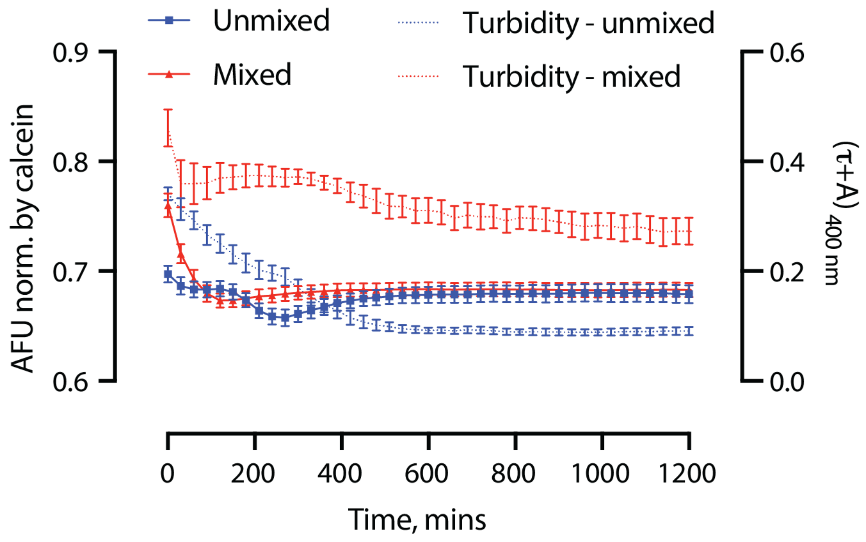


**Fig. S7.** Changes in calcein fluorescence (left y-axis) and turbidity (right y-axis) over the course of a generation 0 incubation comparing samples resuspended and not resuspended at the beginning. Measurements were done every 30 mins. n=10, error bars are SE.

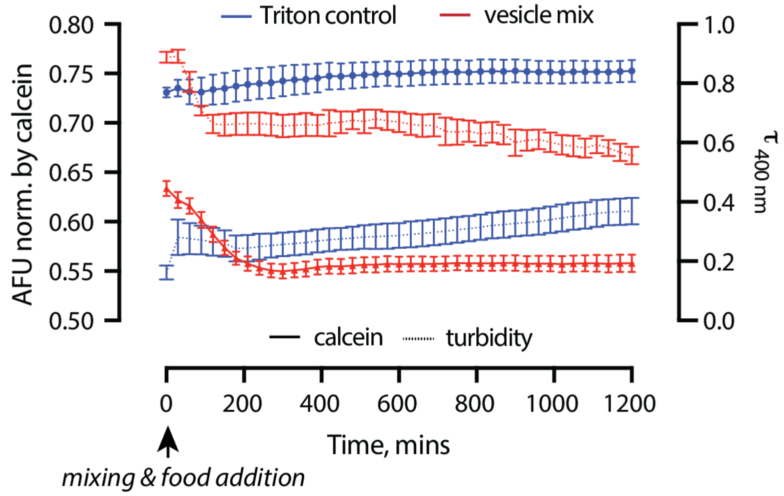


**Fig. S8** Change in calcein fluorescence and turbidity over the course of a generation 1 incubation. Measurements were done every 30 mins. n=10, error bars are SE.

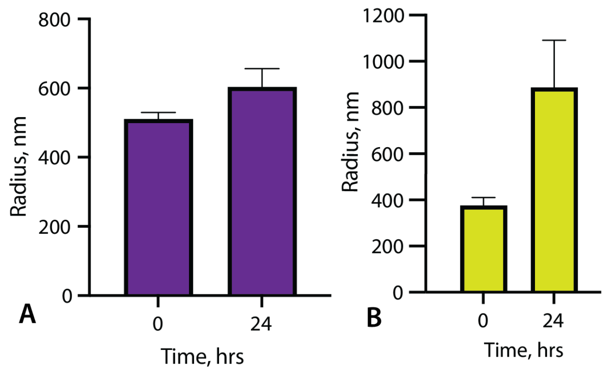


**Fig. S9.** Changes in vesicle size measured by DLS over the course of a generation 0 (**A**) and generation 1 (**B**) incubations. n=8, error bars are SE.

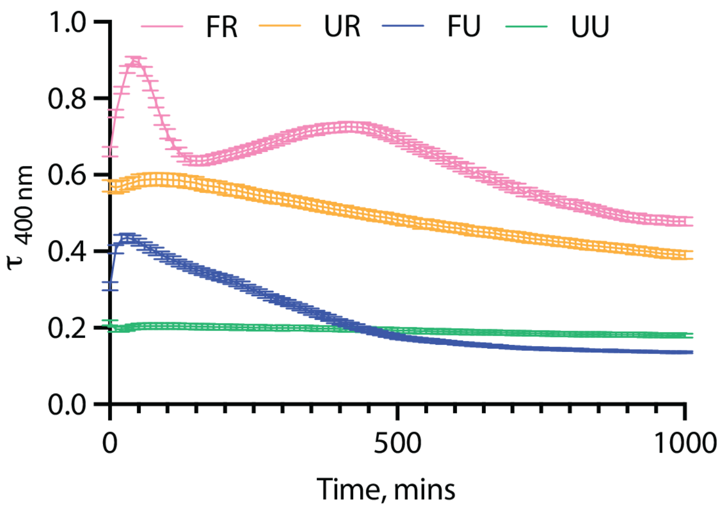


**Fig. S10.** Turbidity timecourses over the first 1000 mins of incubation for the generation 1 of the four experiment types. Measurements were done every 10 mins. n=48 for all but FR, for FR n=96, error bars are SE.

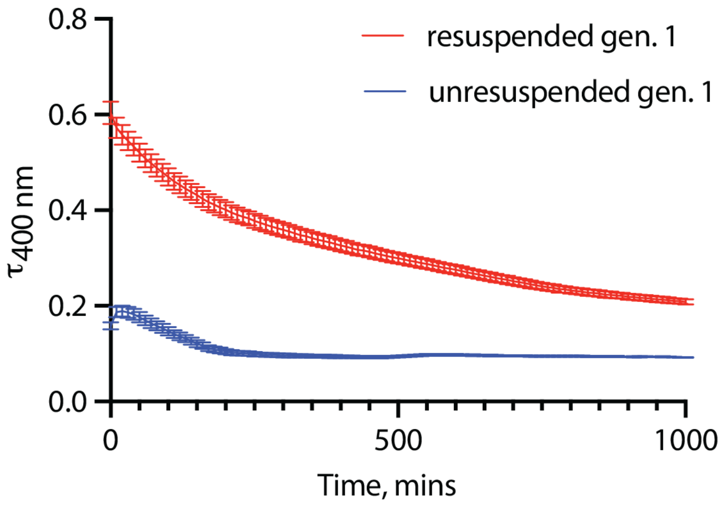


**Fig. S11.** Turbidity timecourses over the first 1000 mins of incubation for a transfer of generation 0 solutions into buffer without food addition with and without resuspending. Measurements were done every 10 mins. n=48, error bars are SE.
